# YfgH is a transmembrane glycine-zipper containing lipoprotein that stabilizes excess cardiolipin and outer membrane proteins during envelope stress

**DOI:** 10.64898/2025.12.22.696046

**Authors:** Ethan Shaigec, Timothy H.S. Cho, Valeria Tsviklist, Justin Bishop, Tracy L. Raivio

**Affiliations:** Department of Biological Sciences, University of Alberta, Edmonton, Alberta, Canada

## Abstract

The outer membrane (OM) of *Escherichia coli* is an essential, asymmetric bilayer composed of lipopolysaccharide and phospholipids that protects cells from environmental stress. While several systems maintain OM integrity during stress, the roles of many outer membrane lipoproteins are less well characterized. Here, we identify YfgH, a glycine-zipper domain containing lipoprotein, as a novel OM-stabilizing factor that functions distinctly from the previously studied SlyB. Deletion of *yfgH* increases sensitivity to detergents and metal chelators, enhances OM permeability, and leads to global loss of OM proteins under stress. YfgH is essential for survival during CRISPR interference-mediated depletion of OM biogenesis genes, and its overexpression rescues these phenotypes. AlphaFold modeling predicts YfgH forms oligomeric ring-like structures similar to SlyB, but genetic suppressor screens indicate distinct mechanisms: *clsA* (cardiolipin synthase) mutations suppress *yfgH* deletion, while *slyB* suppressors map to the Mla lipid transport pathway. Biochemical analyses reveal YfgH interacts with multiple OM proteins, especially under stress. Together these findings suggest YfgH functions by stabilizing cardiolipin-rich nanodomains, representing a novel response to OM stress. This work expands our understanding of glycine-zipper lipoproteins as stress-specific membrane stabilizers and highlights the critical role of YfgH in preserving OM integrity during envelope perturbation.

**Importance:** The bacterial outer membrane is essential for survival in harsh environments, yet how Gram-negative cells maintain its stability under stress remains less understood. We identify YfgH, a previously uncharacterized outer membrane lipoprotein, as a key factor that preserves membrane integrity during envelope stress. Our findings show that YfgH stabilizes cardiolipin-rich nanodomains and protects outer membrane proteins when biogenesis pathways are compromised. This work reveals a new mechanism of outer membrane maintenance and expands the functional repertoire of glycine zipper– containing lipoproteins, highlighting a broader family of stress-specific membrane stabilizers that may be conserved across Gram-negative bacteria.

## Introduction

Gram-negative bacteria are defined by a tri-layered envelope that is critical for survival in harsh environments [1,2]. In most diderm bacteria such as the model organism *Escherichia coli* (*E. coli*), the outer membrane (OM) is an asymmetrical bilayer composed mostly of glycerophospholipids (GPL), lipopolysaccharide (LPS) and proteins [1,3]. LPS resides in the outer leaflet and tightly packs together through salt bridges formed between phosphate groups, conferring the OM with decreased permeability [4].

The inner leaflet of this layer is composed mostly of three different GPL: phosphatidylethanolamine (PE), cardiolipin (CL), and phosphatidylglycerol (PG) [5,6]. The outer membrane proteome on the other hand can be divided into outer membrane proteins (OMPs) and lipoproteins [1,2]. OMPs comprise β-barrel structures integrated in the OM and can carry out a variety of functions including acting as porins and non-covalent binding of the cell wall. Recent work by atomic force microscopy suggests that OMPs and LPS molecules undergo phase separation in which they form distinct islands of like molecules, suggesting that the OM is a highly organized structure [6].

Lipoproteins on the other hand, are lipidated at an N-terminal cysteine residue allowing them to anchor themselves to the OM [1]. The functions of many OM lipoproteins are uncharacterized; however, previous studies have shown that some can form complex structures in the OM and take part in a wide variety of tasks from envelope biogenesis to signaling [7,8,9].

The proteins and lipid components of the OM cannot be synthesized on site; rather, they are manufactured in the cytoplasm or at the inner membrane (IM) and then transported to the OM [10,11,12]. Due to this topology problem, these components must be trafficked to and inserted into the OM without disrupting OM stability [2]. LPS biogenesis can be separated into two parts: synthesis of the mature LPS molecule and its trafficking to the outer membrane [13]. The synthesis of LPS occurs through the Ratz pathway, which takes place in the cytosol and IM [4,13,14]. Mature LPS molecules ready for transport interact with the Lpt machinery for transport to the OM [14,15]. The Lpt transport system is a trans-envelope structure composed of 7 different proteins bridging the IM to the OM. LPS is extracted out of the IM by the ABC transporter LptBCFG, where it is then trafficked across the periplasm by the continuous bridge formed by LptA [15]. A complex of an OM integral OMP LptD and lipoprotein LptE allow insertion of the LPS molecule into the OM [16]. Similarly, to LPS synthesis, biogenesis of OMPs also begins in the cytoplasm, where, after secretion through the SecYEG translocon, the nascent unfolded OMP is trafficked by periplasmic chaperones such as Skp, SurA and DegP to the Bam complex for insertion into the OM [17].

Given the complexity and essentiality of these pathways, being able to respond to perturbations to these biogenesis pathways or to other OM stresses that disrupt this asymmetrical layer is essential to cell survival [18]. To maintain asymmetry, *E. coli* uses diverse systems such as the Mla (maintenance of lipid asymmetry) system and phospholipase PldA [19,20]. In the event of partial loss of asymmetry due to stress and mislocalization of GPL to the outer leaflet, the Mla system composed of MlaBCDEF transports mislocalized GPL in the outer leaflet of the OM back to the IM [20]. In concert with this, the phospholipase PldA degrades mislocalized GPL, effectively removing them from the outer leaflet [21]. Along with these systems, the cell also employs different stress responses such as the Cpx, Rcs, PhoPQ and σ^E^ systems to sense and alleviate envelope stress by changing the expression levels of their respective regulons [18,22]. However, the molecular mechanisms behind how many stress response regulon members combat stress remain elusive.

One such example is the putative lipoprotein YfgH. Previous work has tied YfgH to OM stability during disruption to OM biogenesis; deletion of *yfgH* during CRISPRi knockdown of LptD decreased cell growth and protein levels of OMPs compared to LptD knockdown in WT background [23]. Additionally, a single mutant of *yfgH* showed increased blebbing and OM shedding during vancomycin treatment. However, the mechanism by which YfgH stabilizes the outer membrane remains elusive. In this study we provide evidence that YfgH is a member of a novel class of envelope stabilizing outer membrane lipoproteins.

## Results and Discussion

### Deletion of the OM lipoprotein *yfgH* leads to increased sensitivity to OM stress and OM permeability

Given that YfgH seems to be involved in envelope stress response, we wished to investigate the sub-envelope localization of YfgH [23]. To address this, we expressed a His-tagged YfgH by leaky expression from plasmid pTrc99A and seperated IM and OM fractions via a sucrose density gradient. Inner membrane and outer membrane fractions were determined by blotting for CpxA and BamA respectively.

YfgH-His was predominately found in the fractions containing the OM marker BamA, suggesting OM localization of YfgH **(Figure 1A)**. The N-terminal region of YfgH contains a cysteine similar to other OM lipoproteins **(Figure 1B)**, and SignalP 6.0 predicted YfgH to be a lipoprotein **(Figure S1)** [24].

**Figure 1.**
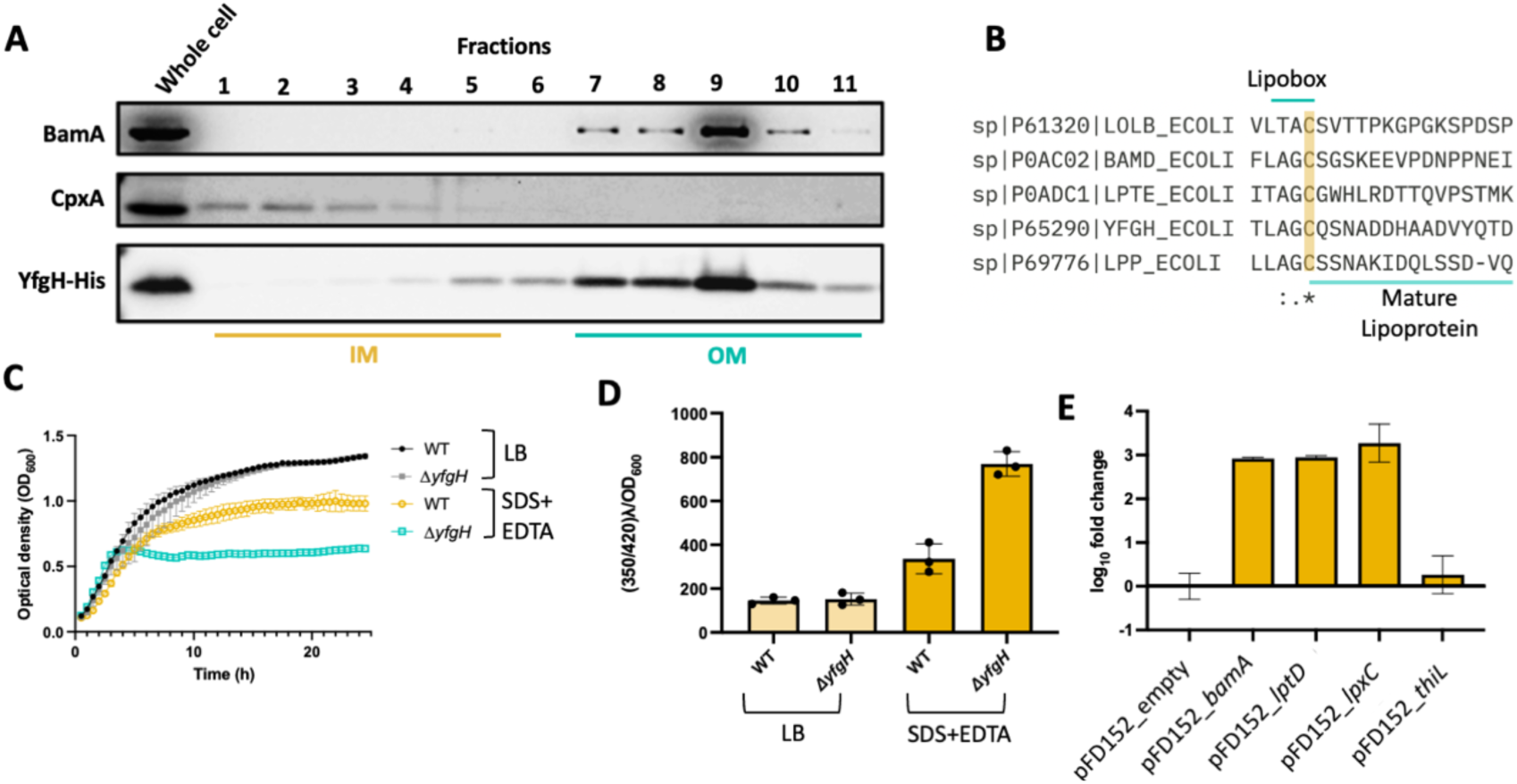
YfgH is a stress regulated Outer Membrane localized lipoprotein involved in OM stability. **(A)** Outer membrane and inner membrane fractionation by sucrose density gradient in cells carrying His-tagged YfgH. Fractions were visualized through immunoblotting with anti-MBP-CpxA (IM control), anti-BamA (OM control), anti-His primary antibodies. (**B)** Sequences of the N-terminal lipobox (indicated by green line) and conserved cysteine (highlighted yellow) of several representative *E. coli* outer membrane lipoproteins and YfgH. (**C)** Growth curves of WT MP1 and Δ*yfgH* in LB vs 0.1% SDS and 0.1mM EDTA. Data is representative of three independent replicates. (**D)** WT and Δ*yfgH* outer membrane permeability measured by fluorescence of NPN (350/420 λ) under LB vs 0.1% SDS and 0.1mM EDTA conditions, normalized to OD600. Data is presented as mean ± SD of 3 biological replicates. (**E)** log_10_ fold change of *yfgH* transcripts in different CRISPRi backgrounds with inducer (500 ng/mL aTc).

A previous study showed that *yfgH* is essential when the OM is compromised by CRISPR interference of the LPS biogenesis gene *lptD* [23]. Given that YfgH seems to stabilize the OM during LPS depletion, we chose to conduct experiments in a recent mouse commensal isolate of *E. coli*, MP1, in which the sequence of *yfgH* at the protein level is identical to that in K-12 *E. coli* **(Figure S2)** [25]. MP1 produces LPS with a full O-antigen, which is missing in K-12 derivative strains. This strain was chosen because we believe it to be a more representative and robust model of *E. coli* capable of survival in its natural hosts, as compared to commonly used laboratory domesticated strains of K-12. For example, previous studies have used MP1 as a model organism to study the role of prophages during colonization [25,26,27].

Given the previous phenotype concerning the LPS biogenesis gene *lptD* we sought to characterize the effect that the single knockout of *yfgH* has on cell physiology in the presence of OM stress. We investigated growth differences of WT and single mutants of *yfgH* in liquid LB with and without SDS and EDTA **(Figure 1C)**. Here, we observed that the Δ*yfgH* strain possessed a significant growth defect in SDS+EDTA compared to WT **(Figure 1C)**. This observation suggests that YfgH is required to combat OM stress. In support of this phenotype not being due to polar effects we were able to complement this phenotype with leaky expression of YfgH **(Figure S4)**. To further test OM stability and permeability, we quantified the uptake of fluorescent dye 1-N-phenyl-ethylamine (NPN) under both LB and SDS+EDTA conditions in WT and Δ*yfgH* cells **(Figure 1D**). NPN is normally excluded from the membrane; however, a compromised OM allows NPN to bind the hydrophobic tail of phospholipid, resulting in fluorescence. Δ*yfgH* cells displayed higher uptake of NPN than WT cells in the presence of SDS+EDTA, suggesting increased OM permeability of the mutant strain under OM stress **(Figure 1D)**. To further establish YfgH as a stress response envelope lipoprotein we performed qPCR in different CRISPRi knockdowns of genes related to, and unrelated to OM biogenesis. It was observed that YfgH was heavily induced in conditions that disrupted the OM, such as CRISPRi knockdowns of LpxC, LptD, and BamA **(Figure 1E)**. Conversely, YfgH was not induced in the empty vector background or in the knockdown of the non-envelope related essential gene *thiL* **(Figure 1E)** [34,35]. Thus, YfgH appears to be an OM lipoprotein involved in the response to OM stress.

### Deletion of *yfgH* negatively impacts the OM during CRISPRi knockdowns of OM biogenesis genes

Given the synthetically lethal phenotype of deleting *yfgH* in a LptD depletion strain in a K-12 derivative strain of *E. coli*, we sought both to replicate this result in MP1 and investigate whether this phenotype could be replicated during knockdowns of other essential outer membrane biogenesis genes. To test this, we knocked down expression of two major OM biogenesis genes, *lptD* and *bamA*, by CRISPRi in WT and Δ*yfgH* backgrounds and assayed for growth defects in both backgrounds **(Figure 2A)**. BamA is an essential β-barrel OMP that is the primary component of the Bam complex responsible for insertion of OMPs into the OM [28]. Growth curves were carried out in the presence and absence of anhydrotetracycline (aTc), the inducer of the CRISPRi system present on plasmid pFD152, which also encodes guide RNAs (gRNA) targeting *bamA* or *lptD* [29,30]. Because overexpression of the CRISPRi system itself has previously been shown to have detrimental effects on cell growth, we included empty vector controls to show that growth defects are due to OM biogenesis gene depletion rather than toxicity due to the CRISPRi system itself [31].

**Figure 2.**
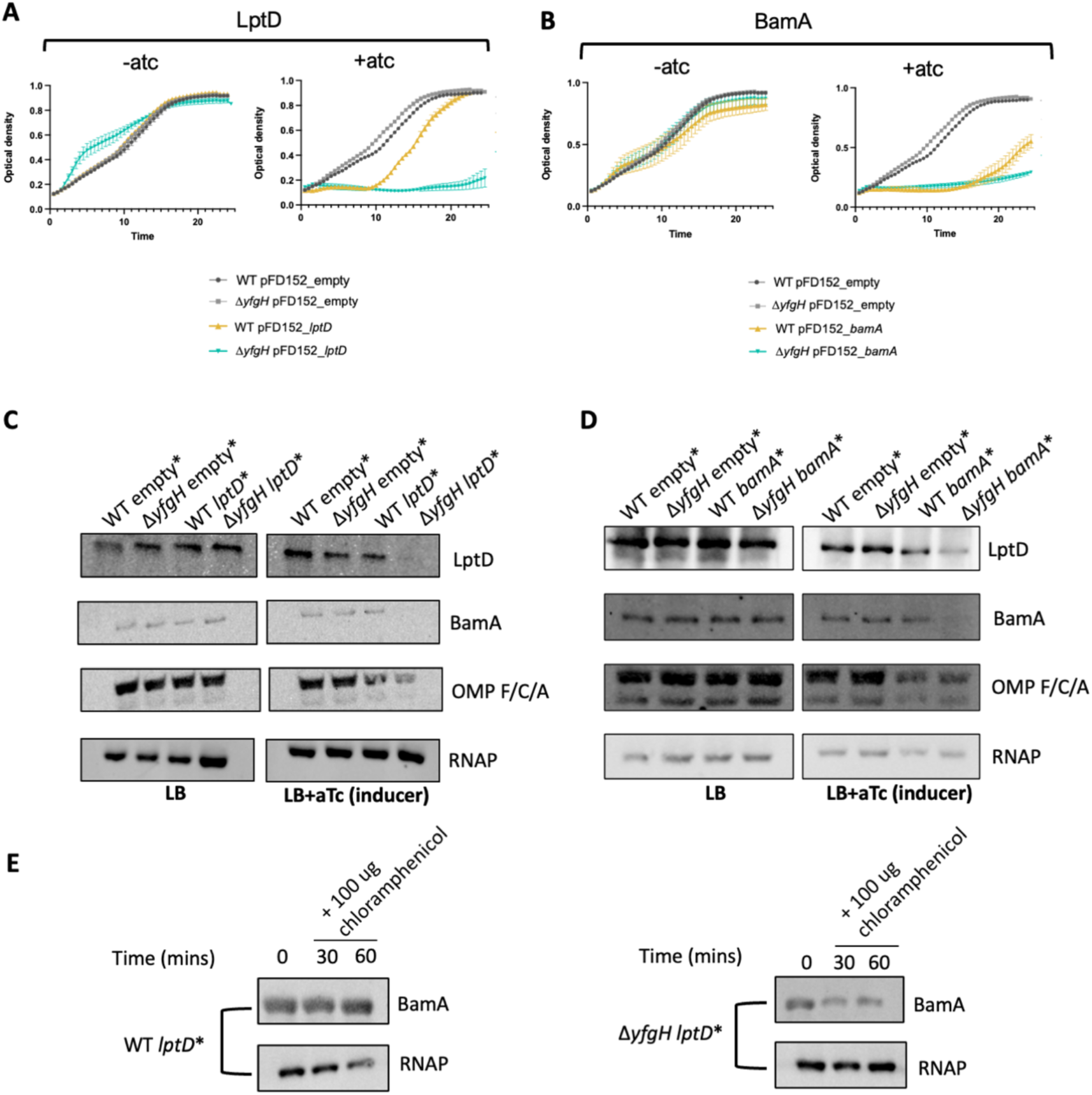
Deletion of *yfgH* shows aggravating interactions with CRISPRi knockdowns of OM biogenesis genes. Growth curves of empty and **(A)** LptD CRISPRi and **(B)** BamA knockdowns in WT vs Δ*yfgH* backgrounds without (left) or with (right) the inducer of the CRISPRi system (500 ng/mL aTc). Data is representative of mean ± SD, with 3 biological replicates. Immunoblots of major OM proteins of cell lysates from CRISPRi knockdown strains of LptD **(C)** and BamA **(D)**. Data is representative of 2 independent replicates. **E** Immunoblots of BamA and loading control (RNAPα) after induction of CRISPRi (500 ng/mL) for 2 hours and then addition of chloramphenicol to inhibit protein synthesis. Data is representative of 2 independent replicates.

Knockdowns of BamA and LptD in a WT background had differing phenotypes; knockdown of BamA seemed to be more lethal than LptD as less growth was observed in the BamA depletion strain at the same concentration of inducer. We believe this to be based on different turnover rates of OMPs vs LPS. Interestingly, little growth was observed when the *yfgH* deletion was introduced into both of these CRISPRi knockdown strains **(Figure 2B)**. Given that LptD and BamA are both OM β-barrels, we investigated if these results could be extended to non-β-barrel proteins involved in OM biogenesis.

For LPS biogenesis we targeted LpxC, the rate limiting enzyme in the synthesis of LPS, and LptA, the periplasmic bridge responsible for transport of LPS to the OM [32]. For OMP biogenesis, we knocked down BamD, the essential lipoprotein of the Bam complex [33]. Knockdowns of LptA, LpxC and BamD in a *ΔyfgH* background grew minimally compared to knockdowns of LptA, LpxC and BamD expression in the WT background **(Figure S5)**. In comparison, no aggravating interactions were observed when knocking down essential genes not directly involved in OM biogenesis, such as *thiL* and *rpoA* as growth kinetics were similar between WT and Δ*yfgH* strains **(Figure S6)** [34,35].

Intriguingly, when knocking down *lolB*, the protein responsible for insertion of OM lipoproteins in the OM, the *yfgH* deletion strain grew better than the WT strain **(Figure S7)**, suggesting that the accumulation of YfgH in the IM may be toxic.

Given the decreased growth of CRISPRi knockdown of OMP and LPS biogenesis in the absence of YfgH, we wished to investigate the OM protein composition of these different strain backgrounds. We probed for the abundance of different OMPs (LptD, BamA, OmpF/C/A) and used RNAPα subunit as a loading control. In absence of aTc, no apparent difference was observed between strain backgrounds **(Figure 2C)**. However, we observed a slight decrease in OM protein levels in the LptD depletion strain when the CRISPRi system was induced. Interestingly, LptD depletion in a *yfgH* deletion background showed severe decrease in OM protein levels. A similar phenotype was observed during knockdown of BamA **(Figure 2D)**.

These results suggested that YfgH may be involved in stabilizing OM components during decreased LPS or OMP biogenesis. Alternatively, YfgH could be involved in changing the transcriptional profile of the cell, specifically by increasing OMP expression to fill in “holes” in the OM. To investigate this, we induced knockdowns of LptD for 2 hours and then used sub-lethal concentrations of chloramphenicol to inhibit any further protein synthesis and assayed for the level of BamA in a WT and Δ*yfgH* background over time. The level of BamA remained fairly stable in the WT background over 30 minutes after the addition of chloramphenicol, whereas we observed loss of BamA in the Δ*yfgH* background, suggesting that YfgH’s role at the OM is not to support transcriptional endogenous stability of these OMPs **(Figure 2E)**. Taken together this suggests that a *yfgH* deletion renders cells less adept at dealing with comprised asymmetry, but also the possibility toxicity arising from incorrect trafficking of the lipoprotein.

### Overexpression of the lipoprotein YfgH rescues CRISPRi knockdowns of major OM biogenesis genes

Given that deletion of *yfgH* renders cells unable to adapt to a compromised OM created by CRISPRi knockdowns, we wondered whether over-supplying the OM with YfgH could improve growth during OM stress. We overexpressed YfgH by leaky expression from the *trc* promoter of vector pTrc99A in knockdowns of BamA and LptD **(Figure 3AB)** [36]. To exaggerate any effects of a potential rescue by YfgH, we induced stronger knockdowns of BamA and LptD by increasing the concentration of aTc used compared to the first set of experiments **(Figure 2AB)**. Cells carrying the pTrc-YfgH vector exhibited improved growth during BamA or LptD knockdown compared to without YfgH overexpression **(Figure 3AB)**. Cells overexpressing YfgH in a BamA knockdown ultimately reached an optical density similar to our empty CRISPRi controls, although showing slower growth rates during exponential phase **(Figure 3B)**. In contrast, over supplying the OM with YfgH in an LptD knockdown background essentially completely rescued its growth defects **(Figure 3A)**. Overexpression of YfgH led to partial rescue of BamA knockdowns around the 10-hour mark, which differed from the rescue of LptD by overexpressing YfgH **(Figure 3A)**.

**Figure 3.**
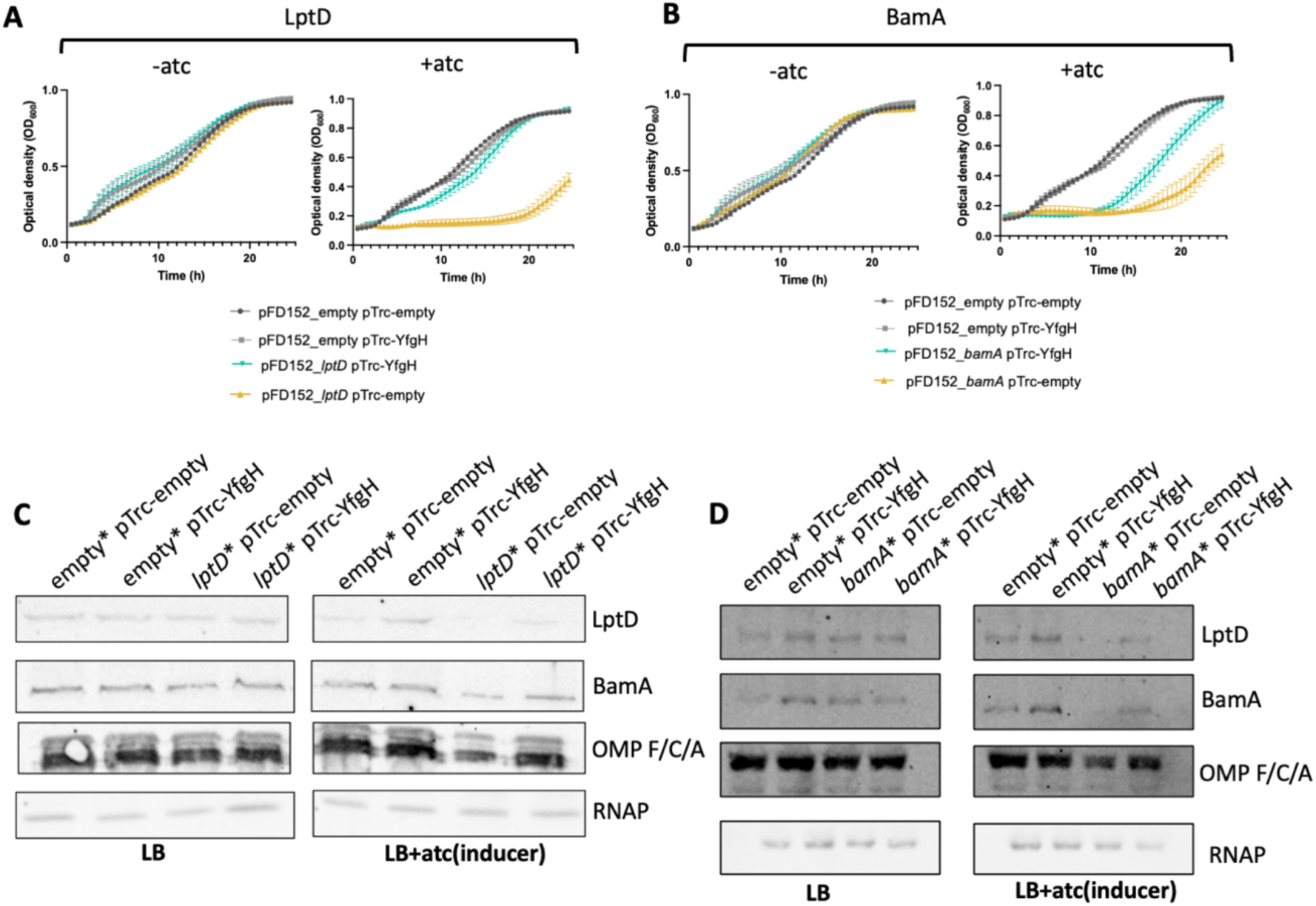
Overexpression of YfgH shows alleviating interactions with CRISPRi knockdowns of OM biogenesis genes. Growth curves of **(A)** LptD and **(B)** BamA CRISPRi knockdowns in WT (pTrc-empty) vs WT YfgH Overexpression (pTrc-YfgH) backgrounds without (left) or with (right) inducer (1000 ng/mL aTc). Data is representative of mean ± SD, with 3 biological replicates. (**B)** Growth curves of empty and BamA Crispri knockdowns in WT (pTrc-empty) vs WT YfgH Overexpression (pTrc-YfgH) backgrounds without (left) or with (right) inducer (aTc). Data is representative of mean +/- SD, where n= 3 biological replicates. (**C, D)** Immunoblots of major OM proteins of cell lysates from above CRISPRi knockdown strains with and without aTc (1000ng/mL). Data is representative of 2 biological replicates.

Similar phenotypes were observed during CRISPRi depletion of the non-β-barrel protein OM biogenesis genes *lpxC*, *lptA*, and *bamD* **(Figure S8)**. In these strains, overexpression of YfgH lead to partial rescue of growth. It is possible that these knockdown recuse differences are due to differences in either half-life of the protein being knocked down or the half-life of the substrates that it is involved in transporting or making. As expected, YfgH overexpression did not lead to any growth changes in CRISPRi knockdown of genes (*thiL* and *rpoA*) not directly related to envelope biogenesis **(Figure S9)**.

Similarly, to the previous assay we investigated OM composition in different strain backgrounds. In the absence of CRISPRi induction, no apparent difference in OMP levels was observed between strains. However, we observed a large global reduction in OMP profile in the LptD and BamA deletion strain. Interestingly, leaky expression of YfgH restored the OM protein profile during LptD depletion **(Figure 3CD)**. A similar phenotype was observed in the case of BamA knockdown and YfgH leaky expression, in which we observed increased outer membrane protein profile in the strain with leaky expression of *yfgH* **(Figure 3D)**. Taken together this suggests that enriching the OM with YfgH renders cells more adept at dealing with comprised asymmetry, which is supported by our previous qPCR results in which we observed a heavy induction of YfgH during OM stress **(Figure 1E)**.

### YfgH is predicted to form oligomeric structures that may function in similar but distinct ways to SlyB oligomers

To further investigate the function of YfgH we created a phylogenetic tree containing all of the known outer membrane lipoproteins in *E. coli* and *yfgH*[37,38,39,]. YfgH clustered in the tree along with other lipoproteins involved in outer membrane stability such as SlyB, ActS, OsmB, Pal, and YiaD **(Figure 4A)** [40,41,42,43]. YfgH is predicted by the Pfam database to contain a two transmembrane (2TM) glycine-zipper domain, which is defined by GXXXG repeats in an alpha helical region that sits within the membrane **(Figure 4B)** [40,45]. SlyB, OsmB, and YiaD are predicted to have this domain as well **(Figure 4B)**. Multiple sequence alignment of the predicted glycine-zipper regions of these proteins confirms the conservation of important GXXXG repeats **(Figure 4C)** [45]. Of particular interest from these OM transmembrane glycine-zippers containing lipoprotein is SlyB, a protein predicted by Alphafold3 (AF3) to be very structurally similar to YfgH **(Figure 4D, F) [**46,47].

**Figure 4.**
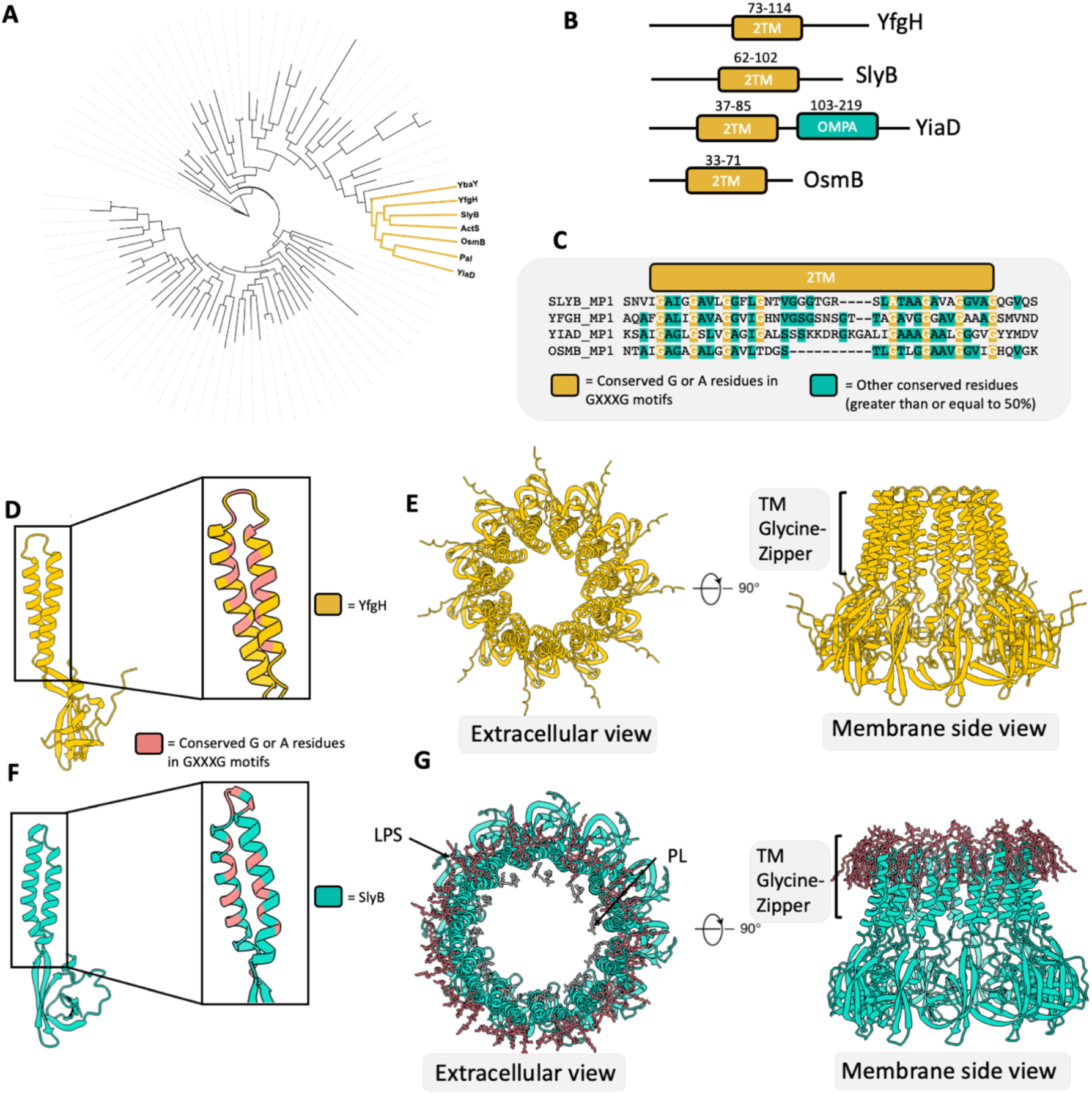
YfgH is member of a larger family of glycine zipper-containing proteins. **(A)** Phylogenetic tree of all known outer membrane lipoproteins in *E. coli*, YfgH is highlighted yellow along with other closely related lipoproteins (**B)** Presence of the transmembrane glycine zipper domain (2TM, yellow) based on panther database and the presence of OmpA C-terminal domain in YiaD (green). Amino acid number is indicated above the protein. **(C)** Multiple sequence alignment of transmembrane Glycine Zipper domain of the four major OM Glycine Zipper containing lipoproteins in *E. coli*. Conserved G and A residues in GXXXG motif are highlighted yellow, whereas other conserved residues are highlighted green. (**D, F)** Alphafold3 structures of YfgH (D) and SlyB (F) with zoomed in image of transmembrane glycine-zipper motif as well as pink highlighted conserved glycine amino acids in zipper domain. € Alphafold3 predicted oligomeric structures of YfgH (YfgH_11_) from membrane view and extracellular view. (**G)** Cryo-EM oligomeric structures of SlyB_11_ forming lipid nanodomains (PL gray, and LPS red) (PDB:70JG)

A previous study suggested that SlyB is involved in responding to loss of lipid asymmetry in the OM [40]. Cryo-EM structures of purified SlyB showed the formation of “lipid nanodomains” within disk-like oligomers of SlyBs when PL islands emerge in the outer leaflet in the OM **(Figure 4G)**. These lipid nanodomains or lipid rafts are formed when SlyB oligomerizes in disk-like shapes around either just mislocalized GPL in the other leaflet or around OMPs surrounded by mislocalized GPL. These lipid nanodomains were thought to break up the emergence of PL islands as a new phase in the OM when LPS is stripped from the outer leaflet (e.g., due to EDTA treatment) and possibly protect encapsulated OM proteins.

Given that YfgH promotes OM stability and is likely structurally similar to SlyB, we wondered if YfgH can adopt a similar disk-like structure to SlyB. To investigate this, we used AF3 to model YfgH homo-oligomers. *yfgH* sequences from *E. coli* strain MP1 with the signal sequence removed and lipidated at the cysteine were used to model higher order YfgH structures using AF3 [48]. The models produced suggested that YfgH could plausibly form disk-like oligomers of various sizes, similar to what was found in the case of SlyB **(Figure 4E) (Figure S10,S11)**. Although we do not present strong biochemical evidence of YfgH oligomers forming, based on similarity to SlyB we predict that this may be a mechanism at play.

### YfgH interacts with a wide range of the IM/OM proteome

The previous interactome and biochemical characterizations of SlyB found that it interacts with various OM proteins. Given the structural and functional similarity of YfgH to SlyB, we investigated if YfgH is similarly able to interact with OM proteins. We used immobilized metal affinity chromatography to purify plasmid-expressed, His×6-tagged YfgH from crude membrane preparations solubilized in β-dodecyl-D-maltoside (DDM). YfgH was purified from cultures grown with or without 5 mM EDTA and in both WT and *clsA::kan* backgrounds. *clsA::kan* is a background that supresses a *yfgH* deletion (**Figure 6**).

Several proteins were affinity purified with YfgH-His×6 as revealed by colloidal Coomassie stain (**Figure 5A**). Interestingly, YfgH appears to co-purify with more proteins, both in terms of number of interacting partners and the amount of protein pulled down, in the presence of 5 mM EDTA compared to the 0 mM EDTA treatment, suggesting that, like SlyB, YfgH may oligomerize more in the presence of envelope stress. Western blotting for known SlyB-interacting proteins confirmed that YfgH interacts with BamA, LptD, and OmpF/C/A **(Figure 5B, S12**). Western blotting also confirmed a slight increase in the amount of BamA and OMPs pulled down relative to the amount of YfgH purified in the presence of EDTA (**Figure 5B, S14**). While we observed a slight decrease in the amount of pull downed proteins in the *clsA::kan* mutant, there was also a proportionate decrease in YfgH purified in this background, suggesting that YfgH can still interact with OM proteins in the absence of the major cardiolipin synthase, although not as efficiently **(Figure 5A)**.

**Figure 5.**
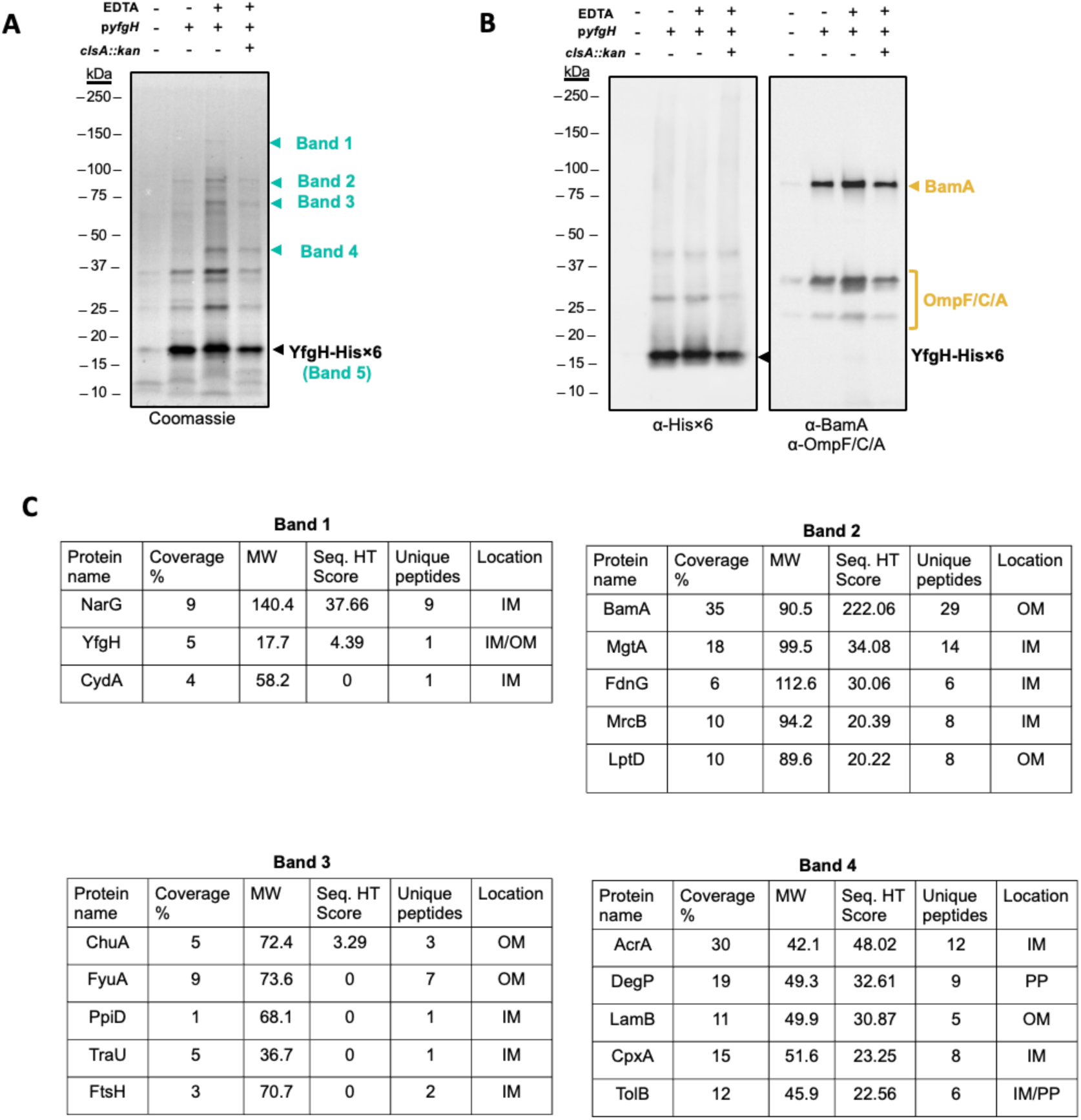
YfgH interacts with OM proteins. **A** Eluted fractions from purification of 6×His-tagged YfgH as visualized using colloidal Coomassie stain. The legend above each image indicates the presence or absence of 5 mM EDTA, the pTrc-YfgH-His×6 or pTrc99A plasmids (p*yfgH* -/+, respectively), and the presence of the *clsA::kan* mutation. **B** Eluted fraction from purification of 6xHis-tagged YfgH western blotted for YfgH-His×6 (α-His×6) or BamA and OmpF/C/A. The legend above each image indicates the presence or absence of 5 mM EDTA, the pTrc-YfgH-His×6 or pTrc99A plasmids (p*yfgH* -/+, respectively), and the presence of the *clsA::kan* mutation. **C** Mass spec results of excised out bands 1 through 4 indicated in panel A. Top 5 membrane associated genes from each band are displayed, along with coverage %, molecular weight (MW), Seq. HT score, Unique peptides, and predicted location.

To further characterize the YfgH interactome, we excised and analyzed five bands (∼140, ∼95, ∼70, ∼45, ∼17 kDa) from the pull-downs in the 5 mM EDTA treatment by mass spectrometry (MS) (**Figure** 5**C****, ST4)**. MS analysis identified the ∼95 kDa band as containing BamA and LptD peptides as well as the ∼17 kDa band containing predominantly YfgH, as expected. Interestingly, the ∼17 kDa band contained peptides that matched the sequence of other small OM lipoproteins including Slp and SlyB, suggesting that YfgH may potentially form heterooligomeric complexes with these proteins. MS analysis of the ∼70 kDa band revealed peptides from FyuA and ShuA, which are OM TonB-dependent receptors for yersiniabactin and heme, respectively, suggesting additional OM protein members of YfgH’s interactome. The most abundant peptides found in the ∼45 kDa band belonged to elongation factor Tu, which may be explained by its very high abundance in the cell [49]. However, peptides corresponding to several envelope proteins were also identified in this band including efflux pump component AcrA, envelope protease and chaperone DegP, and the OM porin LamB. Interestingly, analysis of the ∼140 kDa band revealed peptides belonging to NarG, a member the IM nitrate reductase complex, suggesting that YfgH may be able to form complexes in the IM, at least when overexpressed. While relatively cursory, our analysis suggests that, similarly to SlyB, YfgH forms complexes with several envelope proteins and this interaction protects these proteins during the loss of OM asymmetry.

### *clsA* mutations suppress YfgH deletion sensitivity to SDS+EDTA

To further investigate the function of YfgH, we screened for suppressors of the SDS + EDTA sensitivity of the Δ*yfgH* mutant. We chose this condition, as opposed to sensitivity resulting from diminished expression of an OM biogenesis gene, because we observed that suppressors of CRISPRi knockdowns appear on CRISPRi plasmids at high frequency (data not shown). As expected, we observed decreased growth of the Δ*yfgH* mutant in comparison to the WT in SDS+EDTA **(Figure 6B)**. We plated the Δ*yfgH* mutant at SDS+EDTA concentrations in which the WT parent strain could grow but not the mutant and isolated suppressors able to grow in this condition. After sequencing the genomes isolated from four suppressor colonies, we found that all suppressor mutations mapped to gene, *clsA*, which encodes the major cardiolipin synthase in the cell **(Figure 6A)** [50,51]. One suppressor was a single base pair change resulting in a missense mutation that converted glycine 15 to arginine. Another suppressor mutation also introduced a single base pair change, resulting in a nonsense mutation that converted a tryptophan codon to a stop codon. Another mutation in a different suppressor strain resulted in a frame shift mutation in *clsA* caused by a single base pair deletion. The final suppressor was a 1209 base pair deletion that resulted in partial deletion of both ClsA and the neighbouring gene OppF. All suppressors completely rescued the growth defect of the *yfgH* deletion in SDS+EDTA **(Figure 6B)**. We were able to isolate suppressor mutations in this *clsA* when replicating the original screen **(Figure 6A).** To verify that these suppressors were due to decreased or abolition of ClsA function, we used P1 transduction to introduce the *clsA::kan* mutation from the Keio collection into WT and *yfgH* mutant strains[52]. The Δ*yfgH clsA::kan* strain possessed a similar phenotype to the initial suppressors **(Figure 6C)**, suggesting that the original suppression observed was due to mutations that inactivated ClsA.

**Figure 6.**
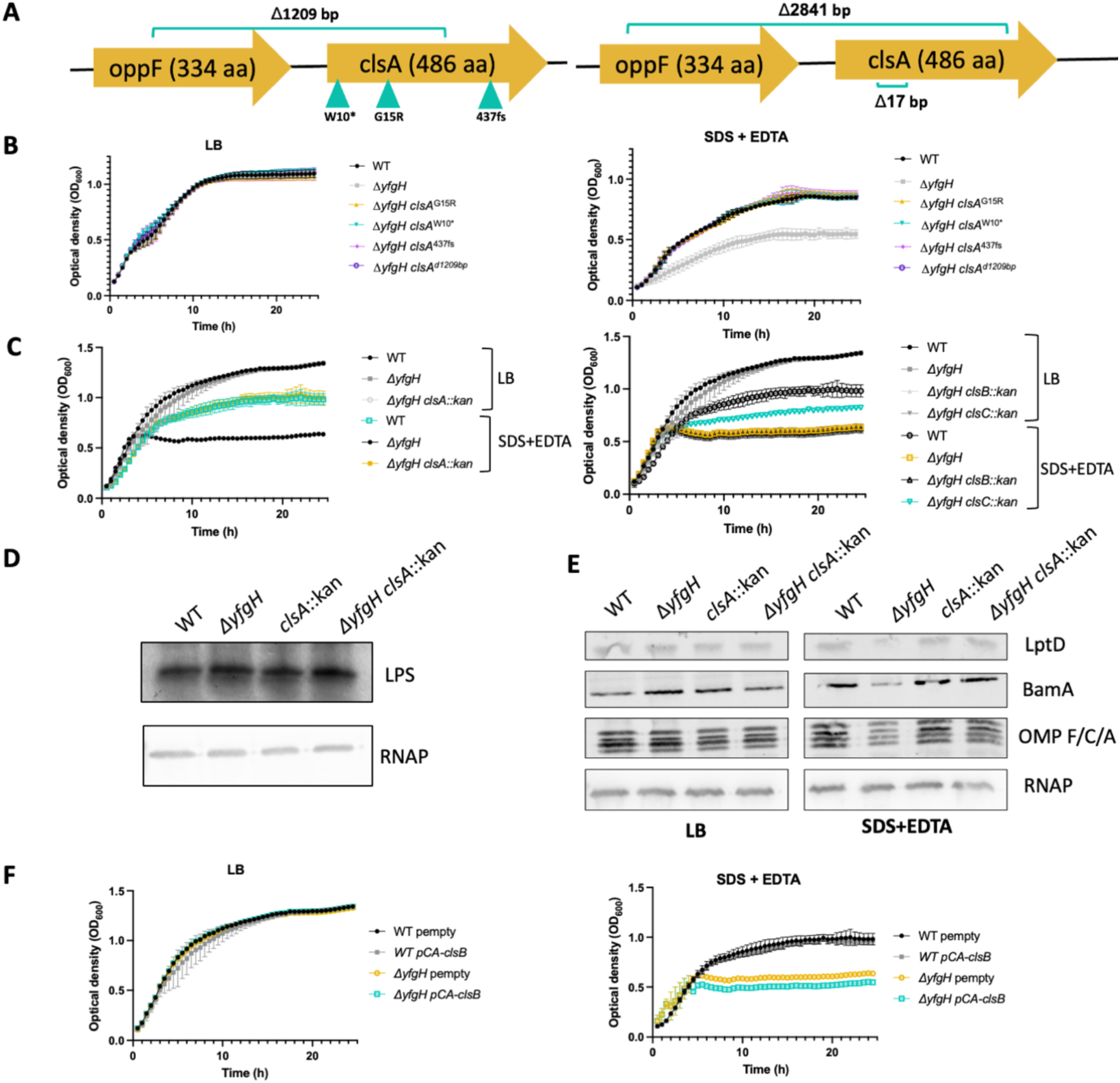
The sensitivity of the *yfgH* mutant to SDS+EDTA is suppressed by deletion of *clsA*. **(A)** Map of isolated from suppressor screen that restore growth of Δ*yfgH* in SDS+EDTA. Each green arrow or bracket indicates different strain sequenced. Left four suppressors are the initial suppressors isolated from single overnight colony. Right two suppressors were isolated from different cultures confirming suppression by mutation in *clsA*. **(B)** Growth curves of WT, Δ*yfgH,* and first 4 isolated suppressors in LB and SDS+EDTA. Data is representative of mean ± SD, with 3 biological replicates. (**C)** Growth curves of WT, Δ*yfgH,*and Δ*yfgH clsA::kan* in LB and SDS+EDTA on the left. Growth curves of WT, Δ*yfgH,* Δ*yfgH clsB::kan,* and Δ*yfgH clsC::kan* in LB and SDS+EDTA on the right. Data is representative of mean ± SD, where n= 3 biological replicates. (**D)** LPS silver stained in different strain backgrounds after SDS-PAGE. Data is representative of 2 biological replicates. **(E)** Immunoblots of major OM proteins of cell lysates from WT, Δ*yfgH, clsA::kan* Δ*yfgH* and *clsA::kan* in LB and SDS+EDTA. Data is representative of 2 biological replicates. (**F)** Growth curves of WT p-empty, Δ*yfgH* p-empty, WT p-ClsB, and Δ*yfgH* p-ClsB in LB and SDS+EDTA on the left. Data is representative of mean ± SD with 3 biological replicates.

*E. coli* expresses two other CL synthases in addition to ClsA: ClsB and ClsC. However, ClsB and ClsC contribute a much smaller proportion of CL in the cell envelope [53]. We observed different phenotypes when deleting the genes encoding these other CL synthases in rescuing the Δ*yfgH* mutant. The Δ*yfgH clsC::kan* double mutant possessed an intermediate level of suppression in comparison to Δ*yfgH clsA::kan*, and a *ΔyfgH clsB::kan* mutant had almost no suppression of the sensitivity to SDS+EDTA at all **(Figure 6C)**. Interestingly, single mutants of the cardiolipin synthases grew on SDS+EDTA normally, suggesting that this phenotype is specific to the absence of YfgH **(Figure S16)**.

Given that SlyB is a similar protein to YfgH and involved in the respose to decreased LPS, we investigated if *clsA* mutations could lead to increased LPS, as CL and LPS are generated from the same initial biogensis pathway [54]. To investigate this, we extracted and silver stained LPS and quantified it against a Western blot of an unrelated loading control protein **(Figure 6D)**. No significant change in LPS was observed between the WT, Δ*yfgH*, *clsA::kan*, and Δ*yfgH clsA::kan* strains. Overall, this suggested that the most likely mechanism of suppression by deleting *clsA* is independent of freeing up more substrate for LPS biogenesis. We next investigated the OM profile to see if *clsA* mutants could suppress the global protein reduction in OM protein during SDS+EDTA treatment **(Figure 6E)**. We observed that mutating *clsA* resulted in a return of OM protein abundance of the Δ*yfgH* strain, similar to amounts in WT. In support of this, we see slight decreased growth in the *yfgH* deletion background when overexpressing *clsB* from a plamsid compared to the *yfgH* deletion background carrying an empty vector **(Figure 6F)** [55,56]. ClsB overexpression was chosen over ClsA overexpression based on that previous studies showed that only over expressing ClsB increases CL in the cells and not ClsA overexpression [55].Taken together, these results suggest that the presence of CL during OM stress without YfgH is detrimental to the cell.

### Sensitivity to SDS+EDTA in a *slyB* mutant is suppressed by deletions in the Mla pathway

To differentiate between the OM stabilizing abilities of SlyB and YfgH, we isolated suppressors to a *slyB* deletion strain via a similar procedure to the screen for *yfgH* deletion suppressors. This allowed us to test if suppressors would arise in *clsA*, like they did for the Δ*yfgH* strain, suggesting a similar mechanism of action between the proteins, or if the suppressors would arise in a different pathway. All four suppressors of the *slyB* mutant’s sensitivy to SDS-EDTA possessed mutations in the Mla pathway that result in decreased or abolished activity of the pathway **(Figure 7A)**. One suppressor was a nine-base pair deletion in *mlaF*. MlaF is part of the inner membrane complex in the Mla pathway that energizes phospholipid transport required for maintaining OM lipid asymmetry[57]. We also found suppressors with a premature a stop codon and a single base pair deletion in *mlaC*., encoding the periplasmic protein responsible for trafficking the PL from the OM back to the IM. The final suppressor had a mutation that mapped to *mlaF*, with an additional mutation in *mltD*, which encodes a protein involved in turnover of peptidoglycan [58]. All suppressors were re-plated on LB agar with and without SDS+EDTA to confirm suppression (Figure **7B-C**).

**Figure 7.**
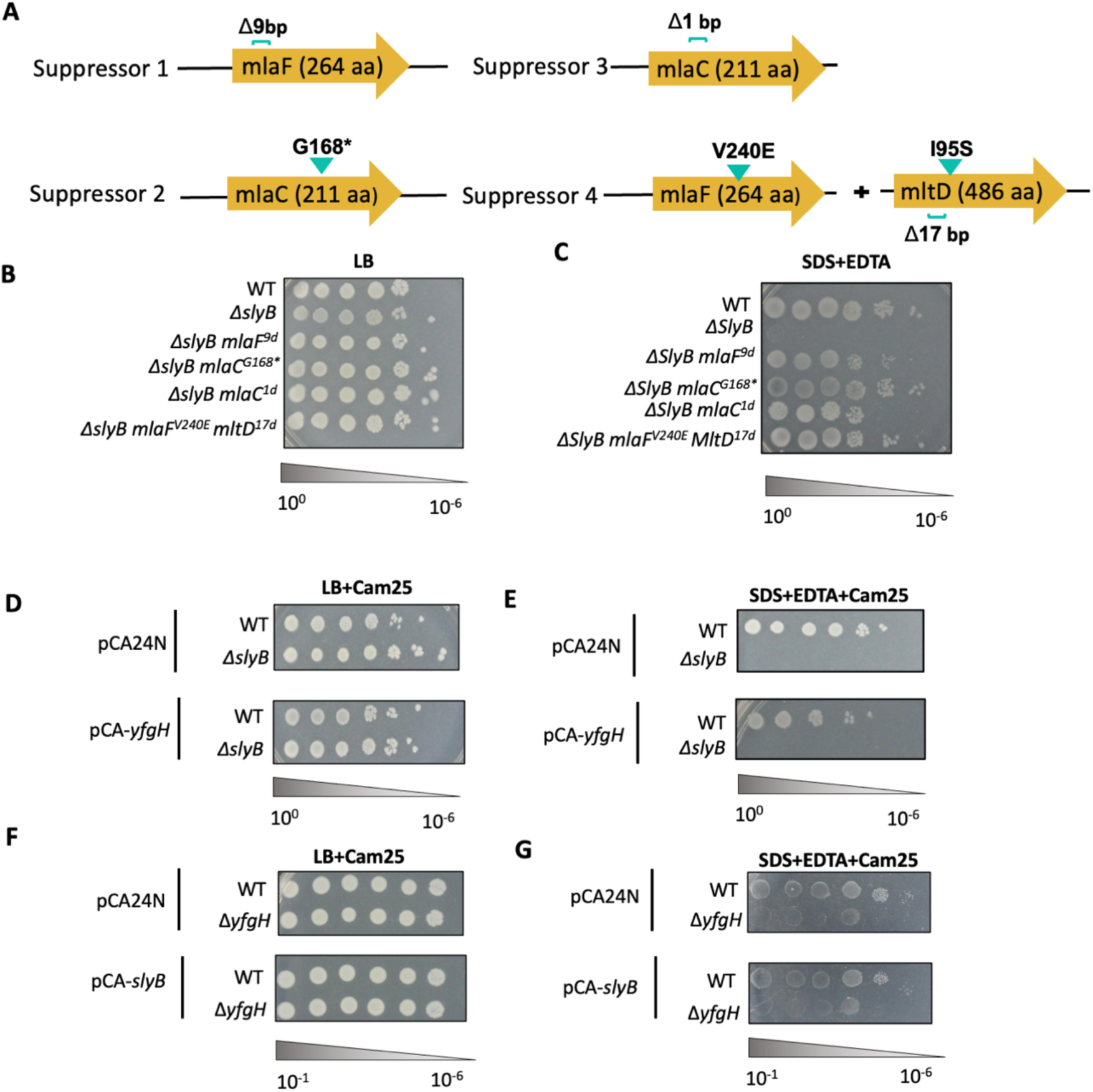
SlyB deletion sensitivity to SDS+EDTA is suppressed by mutation of components of the Mla pathway. **A** Maps of four isolated from suppressor screen to restore growth of Δ*slyB* in SDS+EDTA **B-C** Plating efficiency of WT, Δ*slyB*, and Δ*slyB* suppressors strains on LB and LB + 0.5 mM EDTA and 0.5 % SDS. Data is representative of 3 independent inoculums. 10-fold dilutions are indicated under the images. **D-E** Plating efficiency of WT and Δ*slyB* carrying pCA-YfgH (YfgH overexpression vector) on LB (with chloramphenicol 25 µg/mL) and LB + 0.5 mM EDTA and 0.5 % SDS (with chloramphenicol 25 µg/mL). Data is representative of 3 independent inoculums. 10-fold dilutions are indicated under the images. **F-G** Plating efficiency of WT and Δ*yfgH* carrying pCA-SlyB (SlyB overexpression vector) on LB (with chloramphenicol 25 µg/mL) and LB + 0.5 mM EDTA and 0.5 % SDS (with chloramphenicol 25 µg/mL). Data is representative of 3 independent inoculums. 10-fold dilutions are indicated under the images.

Because suppressors of the Δ*slyB* phenotype arose in a different gene than Δ*yfgH*, we suggest that SlyB stabilizes the OM in a distinct way from YfgH. It should be noted that similar suppressors to global reduction in outer membrane protein profile have rendered similar suppressors, in which a BamD deletion and *bamA* gain of function strain with suppressors that increase PL levels in the OM can suppress lysis of this strain [59]. To further test possible redundancy between YfgH and SlyB we overexpressed YfgH or SlyB by leaky expression form plasmids in a Δ*slyB* or Δ*yfgH* mutant strains and assayed rescue of growth on SDS+EDTA **(Figure 7D-G)**. We observed no growth difference in LB between WT and the Δ*slyB* strain when expressing YfgH **(Figure 7D)**. We also observed that expressing YfgH in the Δ*slyB* strain did not benefit the mutant at all in SDS+EDTA, suggesting the function of these proteins to be distinct **(Figure 7E)**. The inverse was true for the Δ*yfgH* strain when we expressed SlyB, once again suggesting a functional difference between these OM stabilizing proteins **(Figure 7F-G)**.

In further support of this conclusion, we performed qPCR on the other glycine-zipper lipoproteins, in similar conditions that were done for YfgH **(Figure 1E)**. Whereas YfgH was heavily induced under conditions that compromised the OM **(Figure 1E)**, this was not seen for other glycine-zipper lipoproteins (SlyB,YiaD, and OsmB) **(Figure S16)**. These lipoproteins however have much higher expression under normal conditions than that of YfgH, which has been predicted to have as little as 7 copies in the cell under LB conditions [60]. This suggests YfgH may be induced in response to particular stimulus OM stress, such as IM/periplasmic accumulation of LPS or OMPs that could result in problematic CL in the OM.

## Conclusion

Our data suggest that the OM glycine-zipper containing lipoprotein YfgH stabilizes the OM during stress in a similar but distinct way to SlyB, suggesting that it is part of a family of OM-stabilizing glycine-zipper lipoproteins in *E. coli*.

Given that SlyB is thought to oligomerize around PL in the outer membrane during stress, we propose the possibility that YfgH oligomerizes around CL in the outer membrane in an analogous manner **(Figure 8)**. Because reducing the amount of CL in the cell supresses phenotypes of *yfgH* deletion, we hypothesize that having CL in the OM during stress without YfgH would be problematic. Similarly, we suggest that that during loss of asymmetry, YfgH could seal away these cone shaped lipids to prevent blebbing and OM lysis **(Figure 8)**. In further support of this model, several of proteins that affinity purified with YfgH have previously been shown to exist in cardiolipin enriched environments through native mass spectrometry, including NarG, BamA, and LptD [61,62,63].

**Figure 8.**
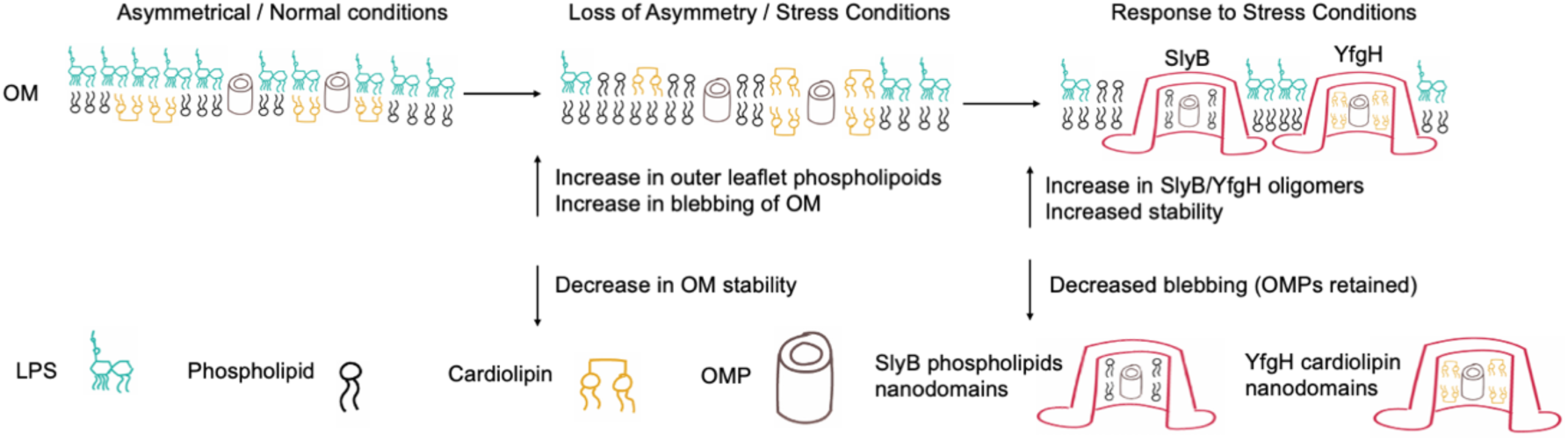
Summary-. Model for YfgH and SlyB OM stabilizing functions. During OM stress cells (such as antibacterial peptides, bile, or EDTA) PL can flip to outer leaflet, which decreases OM stability. YfgH and SlyB cooperate and stabilize different lipids by forming lipid nanodomains around them and as well as encapsulating OMPs. YfgH forms CL nanodomains whereas SlyB forms PL nanodomains.

Overall, our results suggest that YfgH is part of an emerging family of outer membrane stabilizing lipoproteins. Many questions remain including, whether and how much the role of similar outer membrane, glycine zippers containing lipoproteins in *E. coli* may be redundant or distinct. Further the role of the glycine zippers in non-*E. coli* species remain relatively unexplored. Overall, this study highlights the versatility of Gram-negative bacteria in deploying distinct yet complementary mechanisms to maintain envelope integrity under stress.

## Acknowledgements

We thank Dr. Casey Fowler for the use of their centrifuge, Dr. Glen Uhrig for the use of the Beckman Type 70ti rotor and Jack Moore at the Alberta Proteomics and Mass Spectrometry Facility for help with proteomics analysis. We further thank Dr. Ken Rachwalski and Dr. Eric Brown for the CRISPRi knockdown library and Dr. Tom Silhavy for BamA, OmpFCA, and LptD antibodies. We are grateful for help from Dr. Kat Pick, Dr. Randi Guest, and Dr. Brent S. Weber for their help and guidance with suppressor assays and analysis.

## Supplementary Material

Supplementary 1 – contains

Supplementary Figures 1-16 and supplementary tables 1-3

Table 1 = Strains

Table 2 = Plasmids

Table 3 = Primers

Supplementary 2 – contains Supplementary table 4

Table 4 = Mass spec results

Supplementary 2 – contains Supplementary table 5

Table 5 = Raw qPCR

## Materials and Methods

### Bacteria Strains and Growth

All strains used in this study are listed in Table S1. Strains were grown in lysogeny broth (LB; 10 g/L Bacto-Tryptone, 5 g/L Bacto Yeast extract, and 5 g/L sodium chloride) and were supplemented with the appropriate antibiotic for plasmid maintenance (100 ug/mL ampicillin, 100 ug/mL spectinomycin, 25 ug/mL chloramphenicol, 50 ug/mL kanamycin). In the case of liquid cultures, strains were grown at 37°C, shaking at 225 rpm. In the case of solid media, LB was supplemented with agar (15 g/L), and cultures were incubated at 37°C.

### Strain Construction

A list of primers used in this study is listed in Table S2.

Chromosomal knockouts were generated through homologous recombination using Allelic exchange or generalized P1 transduction. Transformations were done by chemical competence.

### Allelic exchange

For strains generated by allelic exchange, 1 kb fragments of sequences upstream and downstream of *yfgH* in MP1 were amplified by PCR. These two fragments were connected via overlap extension PCR to generate the deletion construct. The deletion construct was then cloned into the suicide vector pRE112 using restriction digestion cloning. The cloned plasmid was transformed into *E. coli* PIR2, extracted by mini-prep and sequenced at the Molecular Biology Service Unit at the University of Alberta. The suicide vector was then transformed into MFDλpir and mated with TC635 (MP1) on 0.3 mm DAP. Successful transconjugants were selected on LB+Cam^25^ and incubated overnight. Double crossovers were selected by plating on LB+5% sucrose and incubating for two nights at room temperature. PCR was then used to screen for the correct mutants.

### Generalized P1 Transduction

Kanamycin cassette containing mutants of *clsABC*, *slyB*, and *mlaFC* genes were created by generalized transduction using P1 lysates generated on the Keio collection as described in [51] Mutations were confirmed by PCR. Kanamycin resistance cassette were removed from the *clsA::kan*, *clsB::kan*, *clsC::kan*, and *slyB::kan* strains using Flp recombinase system as described in [51].

### Construction of His-Tagged Protein Overexpression Vectors

YfgH coding regions were amplified from the MC4100 chromosome using PCR using Phusion High-fidelity DNA polymerase (ThermoFisher) and primers containing 17 SacI/XbaI cut sites and a C-terminus 6xHis tag. pTrc99A plasmid DNA and amplified PCR product were double digested with SacI and XbaI Fast Digest restriction endonucleases (Invitrogen) following the manufacturers protocol purified with the QIAquick PCR cleanup kit (Qiagen). Digested PCR product was ligated into digested pTrc99A plasmid using T4 DNA ligase (Invitrogen) following the manufacturer’s protocol and a modified incubation time of 16 hours at 16℃. Ligated plasmid was transformed into One Shot TOP10 chemically competent E. coli (Invitrogen) and plated on LB agar supplemented with 100µg/mL ampicillin. Inserts were verified by PCR using pTrc99A_F and pTrc99A_R primers and DNA sequencing (Table S2). Protein expression in MP1 was verified by SDS-PAGE and western blotting with anti-His antibodies (Figure S3).

### Growth Curves

Growth curves were conducted in 96-well plates at 37°C with shaking at 225 rpm. LB was supplemented with EDTA and/or SDS at various concentrations or anhydrotetracycline in the case of CRISPRi knockdown growth curves. Antibiotics for plasmid maintenance were used as appropriate. The optical density (OD600) was measured every 30 minutes for 24 hours to measure growth. All data was collected as biological triplicates and graphs were plotted with GraphPad PRISM.

### N-phenyl-1-napthyllamie (NPN) OM permeability assay

N-phenyl-1-napthylamie (NPN) was used to quantify OM permeability in different strains in the presence of sublethal concentrations OM damaging agents (EDTA + SDS). Strains were grown overnight in biological triplicates and then sub-cultured and grown until 0.5 OD600. Cells were then washed in NPN buffer (5mM HEPES, 5mM glucose, pH 7.4) and then resuspended in the same buffer. 200uL of each biological triplicate was then transferred to 96 well plates, and NPN was added to a final concentration of 10 uM. EDTA+SDS was then added, cells were incubated for 20 minutes at 37°C with shaking, and then fluorescence (excitation 350 nm, emission 420 nm) was measured. Graphed values show values of fluorescence standardized against OD_600_.

### SDS-PAGE and Western Blotting

Cultures were grown in the same conditions as for other experiments. Cells were collected and standardized to an OD of 1.0 in 100 µL Cells were washed three times in phosphate buffered saline (PBS) and resuspended in water and 2× Laemmli buffer (Sigma). Samples were heated at 95°C for 5 minutes and cooled to room temperature before being separated on 10% or 4-15% tris-glycine SDS-PAGE gels and transferred to a nitrocellulose or PVDF membranes using a semi-dry transfer (BioRad). Membranes were blocked in 5% skim milk dissolved in Tris-buffered saline with 0.1% Tween-20 (TBST) for overnight at 4 degrees Celsius. Membranes were then incubated with primary antibodies (rabbit anti-LptD, anti-BamA, anti-OmpFCA or mouse anti-RNAPα (BioLegend) or His-antibody antibodies) in 2% bovine serum albumin (BSA) in TBST for 30 minutes. Membranes were washed four times with TBST (5 minutes each time) and incubated with fluorescent IRDye680RD (goat anti-mouse) and/or IRDye800 (goat anti-rabbit) secondary antibodies in 5% milk in TBST for 30 minutes. Membranes were then washed again with TBST four times and then imaged using a Bio-Rad ChemiDoc imager.

### Efficiency of Plating

E. coli strains were grown overnight in 5 mL of LB at 37 degrees Celsius with agitation. The culture was diluted to an OD= 0.1 and grown to mid-log phase at 37 °C with agitation and standardized to 0.5 OD. Serial dilutions were then plated onto LB or LB + SDS and EDTA media and incubated overnight at 37°C.

### Suppressor Screen

Overnight colonies of Δ*yfgH* or Δ*slyB* were saved as cryo-stocks to be used as parent strains for identification of SNPs. Overnight cultures were also plated on LB agar containing 1.5% SDS and 1.5 mM EDTA at 37 °C for the Δ*yfgH* strain and 0.75% SDS + 0.75 mM EDTA at 37 °C for the Δ*slyB* strain and incubated until colonies formed. Cryo-stocks of the suppressors were then made, and suppression was confirmed by phenotypic assays. Genomic DNA of the parent and suppressors was then prepped using the Lucigen MasterPure Complete DNA and RNA purification kit, and concentration and quality of gDNA was confirmed via NanoDrop 2000c.

### Genome sequencing

Pure cultures were grown by streaking from glycerol cryo-stocks and then picking single colonies and growing overnight. DNA was extracted from overnight cultures using the Lucigen MasterPure Complete DNA and RNA purification kit, and the concentration and quality of genomic DNA (gDNA) was checked using a NanoDrop 2000c. Library preparation and whole-genome sequencing were performed by the Microbial Genome Sequencing Centre (MiGS, Pittsburgh, PA). Libraries were prepared with the Illumina Nextera kit and sequenced using the NextSeq 550 platform.

To identify mutations in suppressors, each colony was sent for whole-genome sequencing at MiGS, as above. SNPs were identified using BreSeq [64] through Galaxy [65], using a parent strain genome as a reference [66].

### Membrane Fractionation by Sucrose Density Gradients

Cellular localization of YfgH was determined by sucrose density gradients and western blotting. Overnight cultures of strains carrying 6xHis-YfgH overexpression vector was subcultured 1:100 in 250mL LB supplemented with 100µg/mL ampicillin and grown to an OD600 = 1.0 at 37℃ with 180 rpm shaking. Cells were pelleted by centrifugation. The pellet was washed in 50mL 10mM Tris-HCl pH 7.5 and re-pelleted. Cells were resuspended in 10mL Tris-Sucrose (TS) buffer (0.75M sucrose in 10mM Tris-HCl pH 7.5) with 50µg/mL lysozyme and 2mM PMSF. 20mL of 1.65 mM EDTA was slowly added and cell suspensions were incubated for at least 10min on ice. Cell membranes were lysed using 2 passes through a French Pressure Cell Press at 20,000 PSI. 1mL of each sample was collected as a whole cell lysate. Cell debris was removed by centrifugation for 20min at 4℃. Total membranes were pelleted by ultracentrifugation at 38,000 rpm for 45min at 4℃ using a 50.2 Ti rotor (Beckman). Membranes were resuspended in 1mL TES buffer (1 vol TS buffer to 2 vol 1.65mM EDTA) by agitation with sterile plastic loops and pipetting up and down.

Membranes were again pelleted by ultracentrifugation as described above and resuspended in 500µL of 25% (w/w) sucrose in 5mM EDTA. 100µL of sample was collected at this point as a total membrane fraction. Membranes were separated using a six-step sucrose gradient (35-65% sucrose (w/w) in 5mM EDTA) and ultracentrifugation. 4mL of each solution was carefully layered in ultracentrifuge tubes and 400µL of membrane was added on top. Membranes were ultracentrifuged for 18hrs at 40,000rpm at 4℃ using a 50.2 Ti rotor (Beckman). 2mL fractions for a total of 12-13 fractions were pipetted off the top of each gradient and stored in 2mL cryotubes at -80℃. 30µL of each fraction was prepared with 30µL Laemmli buffer and visualized with SDS-PAGE and western blotting. Inner and outer membrane fractions were identified by CpxA and BamA localization respectively.

### LPS Silver Staining

Overnight cultures of strains of interest were pelleted and washed twice with PBS and adjusted to a final OD of 1.0 in 1 mL. They were then resuspended in 50 uL of lysis buffer (2% SDS, 4% 2-mercaptoethanol, 10% glycerol, 1M Tris pH 6.8, bromophenol blue) and incubated at 95℃ for 10 minutes. Each smaple was split into two aliquots, one of which was used to Western blot for RNAP (loading control), and the other was used to silver stain for LPS. For LPS gels, 10 uL of 2.5 mg/mL of proteinase K was added to the sample and incubated the samples at 56℃ for 1 hour. Samples were then run on 12% SDS-PAGE gel (with 19:1 acrylamide/bis-acrylamide) with a 4 % stacking gel. After separation, the gel was fixed with 8.4 mL EtOH, 1 mL acetic acid, 10.6 mL dH_2_O overnight and oxidated with solution of 8.4mL EtOH, 1mL acetic acid, 1.4 mL 10% sodium periodate, and 9.6 mL of dH_2_O for 5 minutes. Gel was then washed with distilled H_2_O 3 times for ten minutes each wash. Gels were then imaged using an iPhone against a white background.

### Chloramphenicol OM protein stability assay

Overnight cultures of CRISPRi knockdown strains were subcultured 1/50 and grown to 0.5 OD in LB+spectinomycin to maintain pFD152. Cultures were then induced with 500 ng/mL of atc to induce the CRISPRi. After allowing to grow for 2 hours the cultures were spun down and washed twice with PBS. Cultures were then resuspended in LB + 100 ug/mL of Chloramphenicol to inhibit protein synthesis. Cells were spun down at set time points (0, 0.5-hour, 1 hour). Timepoints were then western blotted using BamA antibodies to probe stability of this protein in the outer membrane and RNAP as loading control.

### Purification of YfgH-6×His

Expression and purification of His×6-tagged YfgH was conducted in a similar manner to other studies [40,67]. Strains were subcultured 1/50 from overnight cultures into 250 ml LB cultures with 100 µg/ml ampicillin and 0.1 mM IPTG and incubated at 37°C with shaking. 5 mM EDTA was added to appropriate cultures after 1 hour of incubation and cultures were grown for an additional 6 hours.

Cells were collected by centrifugation at 5,700 RPM for 20 minutes and resuspended in 30 ml of Tris buffer (20 mM Tris-HCl pH 8). Cells were pelleted again, supernatants were aspirated, and pellets were left to freeze overnight at -80°C.

The next day, pellets were thawed and resuspended in 22 ml of Tris buffer with one tablet of Roche cOmplete EDTA-free Protease Inhibitor Cocktail and 100 µl of 5 mg/ml lysozyme. Cells were lysed using three passes in a French press and cell lysates were clarified with a spin at 20,000×g for 20 minutes at 4°C. Clarified lysates were transferred to a clean ultracentrifuge tube and spun for 1 hour at 38,000 RPM at 4°C in a Beckman Type 70ti rotor to pellete crude membrane fractions. Supernatants were aspirated and saved and pellets were carefully dislodged and resuspended in about 500 µl of membrane solubilization buffer (50 mM Tris-HCl pH 8, 150 mM NaCl, 20 mM imidazole, 1% n-dodecyl-β-D-maltopyranoside [DDM], 1 cOmplete PIC tablet per 10 ml of buffer). Membrane pellets were allowed to solubilize in buffer for about 1 hour at 4°C with gentle rocking. Insoluble precipitates were observed in some samples and were removed by centrifugation. A small aliquot of each crude membrane preparation was saved for protein quantification by BCA assay (Pierce) and subsequent SDS-PAGE analysis.

Purification of YfgH-His×6 was conducted using HisPur Cobalt Resin (Thermo Scientific). Briefly, 1 ml of slurry was added to a gravity chromatography column, solution was allowed to flow-through, and resin was equilibrated with 2 bed volumes of wash buffer (50 mM Tris-HCl pH 8, 150 mM NaCl, 50 mM imidazole, 0.03% DDM). After wash buffer was allowed to flow-through, crude membrane preparations were added to the resin along with an addition 1-2 ml of wash buffer. Columns were capped and sealed, and binding was allowed to proceed for 1 hour at 4°C. Unbound proteins were allowed to flow-through and were collected. Resin was washed twice with 5 ml (10 bed volumes) of wash buffer, with each wash being collected and prepared for SDS-PAGE. Samples were eluted twice with 750 µl of elution buffer (50 mM Tris-HCl pH 8, 150 mM NaCl, 500 mM imidazole, 0.03% DDM). All collected samples (input/crude membrane preps, flow-through, washes, and elutions) were processed for SDS-PAGE and colloidal Coomassie staining for total protein visualization and Western blotting for identification of specific proteins.

### Mass Spectrometry

Mass spectrometry was carried out at the Alberta Proteomics and Mass Spectrometry Facility. In-gel samples were reduced (10mm BME in 50mm bicarbonate), alkylated (55mM iodoacetamide in 50mm bicarbonate), and digested overnight at 37C (Promega sequencing grade modified trypsin). Tryptic peptides were extracted from the gel twice with 69% water, 30% acetonitrile, 1% formic acid.

Peptides were separated and analysed using a Vanquish Neo UHPLC system (Thermo Scientific) and an EASY-Spray capillary HPLC column (ES75150, Thermo Scientific) coupled to an Orbitrap Exploris 480 mass spectrometer (Thermo Scientific). The mass spectrometer was operated in data-dependent acquisition mode with a resolution of 60,000 and m/z range of 350–1700. Multiply charged ions were fragmented via HCD dissociation with an NCE of 28, and spectra of their fragments were recorded in the orbitrap at a resolution of 15,000. Data was processed using Proteome Discoverer (Thermo Scientific, version 3.0) and the database (Uniprot, UP000284592) was searched using SEQUEST (Thermo Scientific). Search parameters included a strict false discovery rate (FDR) of .01, a relaxed FDR of .05, a precursor mass tolerance of 10ppm and a fragment mass tolerance of 0.01Da. Peptides were searched with carbamidomethyl cysteine as a static modification, and oxidized methionine and deamidated glutamine and asparagine as dynamic modifications.

### RNA isolation and cDNA synthesis

Samples used for the RNA extraction were prepared by diluting overnight cultures 1:50 into 2 ml fresh LB containing appropriate concentrations of antibiotics. CRISPRi knockdowns by pFD152 were induced with 1000 ng/mL aTc and grown to early log stage (OD_600_ of ∼0.35 to 0.4) at 37°C. Then, 1 mL samples were harvested, and RNA was isolated using a QuickExtract nucleic acid extraction kit as per manufacturers’ protocol (LGC Biosearch Technologies). Isolated RNA was resuspended in 35μL of nuclease-free water (IDT) and 1μL of RiboGuard RNase inhibitor (LGC Biosearch Technologies) to prevent RNA degradation. Quality control of the extracted RNA was performed using an Agilent 2100 Bioanalyzer System at the University of Alberta Molecular Biology Service Unit (MBSU).

For cDNA synthesis, RNA concentrations were standardized to 500 ng/μL, and 1μg of RNA was mixed with 3.3μL of 300ng/μl of random primers (Invitrogen), 1μL each of all deoxynucleoside triphosphates (10 mM; Invitrogen), and nuclease-free water to a final volume of 10μL. The random primers and RNA were first allowed to anneal at 70°C for 10 min and then at 25°C for 10 min. After the annealing step, the RNA-primer hybridization mixture was added to 4μL of 5× 1st Strand Buffer, 2μL of 100mM DTT, 1.5μL of 20U/μl SUPERaseIn, 2.5μL of 200U/μl SuperScript II, and nuclease-free water to a final volume of 20μL. The mixture for cDNA synthesis was incubated at 25°C for 10 min, 37°C for 1 h, 42°C for 1 h, and 70°C for 10 min.

### qPCR

The qPCR primers were designed to amplify approximately 75 nucleotides in the last 500bp of each gene of interest and are listed in **Table S2**. qPCR was performed with a 96-well microtiter plate containing a 10μl reaction mixture of the 2× QPCR Mastermix (*Dynamite*), 2.5μl of a 3.2 μM stock solution of each primer and 2.5μl of a 500ng/μl cDNA template. The 2× QPCR Mastermix (*Dynamite*) used in this study is a proprietary mix developed and distributed by the Molecular Biology Service Unit (MBSU), in the Department of Biological Science at the University of Alberta, Edmonton, Alberta, Canada. It contains Tris (pH 8.3), KCl, MgCl2, glycerol, Tween20, DMSO, dNTPs, ROX as a normalizing dye, SYBR Green (Molecular Probes) as the detection dye, and an antibody inhibited Taq polymerase. The qPCR was carried out by incubation at 95°C for 15 s, followed by 60°C for 1 min. This cycle was repeated for a total of 40 times, using a 7500 Fast Real-Time PCR system (Applied Biosystems).

Relative amounts of PCR product were determined by monitoring the number of cycles required to reach a threshold level of fluorescence (cycle threshold [*CT*]) for each gene and subtracting this number from the *CT* value for an endogenous control gene known not to be regulated by envelope stress. We used the *rpoD* gene for this purpose. The resulting delta CTs (*ΔCTs*) were compared between the wildtype strain carrying the pFD152 empty vector and the strains harboring CRISPRi knockdowns of the indicated OM and non-OM biogenesis genes to obtain delta-delta CT values (*ΔΔCTs*), which represented the fold change in gene expression between different strains.

## REFERENCES

1. Nikaido, H. Molecular Basis of Bacterial Outer Membrane Permeability Revisited. Microbiol Mol Biol Rev 67, 593–656 (2003).

2. Silhavy, T. J., Kahne, D. & Walker, S. The Bacterial Cell Envelope. Cold Spring Harbor Perspectives in Biology 2, a000414–a000414 (2010)

3. J. Sun, S. T. Rutherford, T. J. Silhavy, K. Casey Huang, Physical properties of the bacterial outer membrane. Nat. Rev. Microbiol. 20, 236–248 (2022).

4. R. L. Guest, T. J. Silhavy, Cracking outer membrane biogenesis. Biochim. Biophys. Acta BBA - Mol. Cell Res. 1870, 119405 (2023).

5. Lugtenberg, B. Composition and function of the outer membrane of escherichia coli. Trends in Biochemical Sciences 6, 262–266 (1981).

6. Benn, G., et al. Phase separation in the outer membrane of Escherichia coli. Proc. Natl. Acad. Sci. U.S.A. 118, e2112237118 (2021).

7. May, K. L. & Grabowicz, M. Outer Membrane Lipoproteins: Late to the party, but the center of attention. Journal of Bacteriology 207, (2025).

8. Cho, T. H., Wang, J. & Raivio, T. L. NLPE is an OmpA-associated outer membrane sensor of the CPX envelope stress response. Journal of Bacteriology 205, (2023).

9. Malojčić, G. et al. LPTE binds to and alters the physical state of LPS to catalyze its assembly at the cell surface. Proceedings of the National Academy of Sciences 111, 9467–9472 (2014).

10. Simpson, B. W. & Trent, M. S. Pushing the envelope: LPS modifications and their consequences. Nat Rev Microbiol 17, 403–416 (2019)

11. Yeow, J. & Chng, S.-S. Of zones, bridges and chaperones–phospholipid transport in bacterial outer membrane assembly and homeostasis. Microbiology 168, 001177 (2022).

12. Sperandeo, P., Martorana, A. M. & Polissi, A. The lipopolysaccharide transport (Lpt) machinery: A nonconventional transporter for lipopolysaccharide assembly at the outer membrane of Gram-negative bacteria. Journal of Biological Chemistry 292, 17981–17990 (2017).

13. Konovalova, A., Kahne, D. E. & Silhavy, T. J. Outer Membrane Biogenesis. Annu. Rev. Microbiol. 71, 539–556 (2017).

14. Tokuda, H. Biogenesis of Outer Membranes in Gram-Negative Bacteria. Bioscience, Biotechnology, and Biochemistry 73, 465–473 (2009).

15. Bertani, B. & Ruiz, N. Function and Biogenesis of Lipopolysaccharides. EcoSal Plus 8, ecosalplus.ESP-0001-2018 (2018).

16. M. N. Webby et al., Lipids mediate supramolecular outer membrane protein assembly in bacteria. Sci. Adv. 8, eadc9566 (2022).

17. Tokuda, H. Biogenesis of Outer Membranes in Gram-Negative Bacteria. Bioscience, Biotechnology, and Biochemistry 73, 465–473 (2009).

18. Hews, C. L., Cho, T., Rowley, G. & Raivio, T. L. Maintaining Integrity Under Stress: Envelope Stress Response Regulation of Pathogenesis in Gram-Negative Bacteria. Front. Cell. Infect. Microbiol. 9, 313 (2019).

19. May, K. L. & Silhavy, T. J. The *Escherichia coli* phospholipase PldA regulates outer membrane homeostasis via lipid signaling. mBio 9, e00379–18 (2018).

20. Yeow, J. & Chng, S.-S. Of zones, bridges and chaperones–phospholipid transport in bacterial outer membrane assembly and homeostasis. Microbiology 168, 001177 (2022).

21. Shrivastava, R. & Chng, S.-S. Lipid trafficking across the Gram-negative cell envelope. Journal of Biological Chemistry 294, 14175–14184 (2019).

22. Guest, R. L., Lee, M. J., Wang, W. & Silhavy, T. J. A periplasmic phospholipase that maintains outer membrane lipid asymmetry in Pseudomonas aeruginosa. Proc. Natl. Acad. Sci. U.S.A. 120, e2302546120 (2023).

23. Babu, M. et al. Genetic Interaction Maps in Escherichia coli Reveal Functional Crosstalk among Cell Envelope Biogenesis Pathways. PLoS Genet 7, e1002377 (2011).

24. Teufel, F. et al. SIGNALP 6.0 predicts all five types of signal peptides using protein language models. Nature Biotechnology 40, 1023–1025 (2022).

25. Lasaro, M. et al. Escherichia coli isolate for studying colonization of the mouse intestine and its application to two-component signaling knockouts. Journal of Bacteriology 196, 1723–1732 (2014).

26. Pick, K., Stadel, L. & Raivio, T. L. *escherichia coli* phage-inducible chromosomal island aids helper phage replication and represses the locus of enterocyte effacement Pathogenicity Island. The ISME Journal 19, (2025).

27. Pick, K., Ju, T., Willing, B. P. & Raivio, T. L. Isolation and characterization of a novel temperate escherichia coli bacteriophage, kapi1, which modifies the O-antigen and contributes to the competitiveness of its host during colonization of the murine gastrointestinal tract. mBio 13, (2022).

28. Doyle, M. T. & Bernstein, H. D. Bama forms a translocation channel for polypeptide export across the bacterial outer membrane. Molecular Cell 81, (2021).

29. Rachwalski, K. et al. A Mobile CRISPRI collection enables genetic interaction studies for the essential genes of escherichia coli. Cell Reports Methods 4, 100693 (2024).

30. Depardieu, F. & Bikard, D. Gene silencing with CRISPRI in bacteria and optimization of DCAS9 expression levels. Methods 172, 61–75 (2020).

31. Sun, L., Zheng, P., Sun, J., Wendisch, V. F. & Wang, Y. Genome-scale CRISPRI screening: A powerful tool in Engineering Microbiology. Engineering Microbiology 3, 100089 (2023).

32. Shu, S. & Mi, W. Regulatory mechanisms of lipopolysaccharide synthesis in escherichia coli. Nature Communications 13, (2022).

33. Wu, T., et al. Identification of a multicomponent complex required for outer membrane biogenesis in escherichia coli. Cell 121, 235–245 (2005).

34. Leonardi, R. & Roach, P. L. Thiamine biosynthesis in escherichia coli. Journal of Biological Chemistry 279, 17054–17062 (2004).

35. Jafri, S., Urbanowski, M. L. & Stauffer, G. V. A mutation in the RPOA gene encoding the alpha subunit of RNA polymerase that affects mete-metr transcription in escherichia coli. Journal of Bacteriology 177, 524–529 (1995).

36. Amann, E., Ochs, B. & Abel, K.-J. Tightly regulated TAC promoter vectors useful for the expression of unfused and fused proteins in escherichia coli. Gene 69, 301–315 (1988).

37. Edgar, R. C. Muscle: Multiple sequence alignment with high accuracy and high throughput. Nucleic Acids Research 32, 1792–1797 (2004).

38. Guindon, S., Lethiec, F., Duroux, P. & Gascuel, O. PHYML online--a web server for fast maximum likelihood-based phylogenetic inference. Nucleic Acids Research 33, (2005).

39. Letunic, I. & Bork, P. Interactive tree of life (itol) V5: An online tool for phylogenetic tree display and annotation. Nucleic Acids Research 49, (2021).

40. Janssens, A. et al. SlyB encapsulates outer membrane proteins in stress-induced lipid nanodomains. Nature (2023)

41. Gurnani Serrano, C. K., et al. Acts activates peptidoglycan amidases during outer membrane stress in *escherichia coli*. Molecular Microbiology 116, 329–342 (2021).

42. Szczepaniak, J. et al. The lipoprotein pal stabilises the bacterial outer membrane during constriction by a mobilisation-and-capture mechanism. Nature Communications 11, (2020).

43. Tachikawa, T. & Kato, J. Suppression of the Temperature-Sensitive Mutation of the bamDGene Required for the Assembly of Outer Membrane Proteins by Multicopy of the yiaD Gene in Escherichia coli. Bioscience, Biotechnology, and Biochemistry 75, 162–164 (2011).

44. Mi, H., Muruganujan, A., Casagrande, J. T. & Thomas, P. D. Large-scale gene function analysis with the PANTHER classification system. Nat Protoc 8, 1551–1566 (2013)

45. Kim, S., et al. Transmembrane glycine zippers: physiological and pathological roles in membrane proteins. Proc. Natl Acad. Sci. USA 102, 14278–14283 (2005).

46. Jumper, J. et al. Highly accurate protein structure prediction with AlphaFold. Nature 596, 583–589 (2021).

47. Varadi, M. et al. Alphafold protein structure database: Massively expanding the structural coverage of protein-sequence space with high-accuracy models. Nucleic Acids Research 50, (2021).

48. Abramson, J. et al. Accurate structure prediction of biomolecular interactions with alphafold 3. Nature 630, 493–500 (2024).

49. Soufi, B., Krug, K., Harst, A., & Macek, B. (2015). Characterization of the E. coli proteome and its modifications during growth and ethanol stress. Frontiers in Microbiology, 6. 10.3389/fmicb.2015.00103

50. Carranza, G. et al. Cardiolipin plays an essential role in the formation of intracellular membranes in escherichia coli. Biochimica et Biophysica Acta (BBA) - Biomembranes 1859, 1124–1132 (2017).

51. Douglass, M. V., Cléon, F. & Trent, M. S. Cardiolipin aids in lipopolysaccharide transport to the gram-negative outer membrane. Proceedings of the National Academy of Sciences 118, (2021).

52. Baba, T. et al. Construction of *Escherichia coli* K-12 in-frame, single-gene knockout mutants: The keio collection. Molecular Systems Biology 2, (2006).

53. Tan, B. K. et al. Discovery of a cardiolipin synthase utilizing phosphatidylethanolamine and phosphatidylglycerol as substrates. Proceedings of the National Academy of Sciences 109, 16504–16509 (2012).

54. Som, N. & Reddy, M. Cross-talk between phospholipid synthesis and peptidoglycan expansion by a cell wall hydrolase. Proceedings of the National Academy of Sciences 120, (2023).

55. Jeucken, A., Helms, J. B. & Brouwers, J. F. Cardiolipin synthases of escherichia coli have phospholipid class specific phospholipase D activity dependent on endogenous and foreign phospholipids. Biochimica et Biophysica Acta (BBA) - Molecular and Cell Biology of Lipids 1863, 1345–1353 (2018).

56. Kitagawa, M. et al. Complete set of ORF clones of escherichia coli ASKA library (a complete set of E. Coli K-12 ORF archive): Unique Resources for Biological Research. DNA Research 12, 291–299 (2006).

57. Malinverni, J. C. & Silhavy, T. J. An ABC transport system that maintains lipid asymmetry in the gram-negative outer membrane. Proceedings of the National Academy of Sciences 106, 8009–8014 (2009).

58. Kaul, M., Meher, S. K., Nallamotu, K. C. & Reddy, M. Glycan strand cleavage by a lytic transglycosylase, MLTD contributes to the expansion of peptidoglycan in escherichia coli. PLOS Genetics 20, (2024).

59. Mikheyeva, I. V., Sun, J., Huang, K. C. & Silhavy, T. J. Mechanism of outer membrane destabilization by global reduction of protein content. Nat Commun 14, 5715 (2023).

60. Moore, L. R., Caspi, R., Boyd, D., Berkmen, M., Mackie, A., Paley, S., & Karp, P. D. (2024). Revisiting the Y-ome of *escherichia coli*. Nucleic Acids Research, 52(20), 12201–12207. 10.1093/nar/gkae857

61. Arias-Cartin, R. et al. Cardiolipin-based respiratory complex activation in bacteria. Proceedings of the National Academy of Sciences 108, 7781–7786 (2011).

62. Chorev, D. S. et al. Protein assemblies ejected directly from native membranes yield complexes for mass spectrometry. Science 362, 829–834 (2018).

63. Fiorentino, F. et al. Dynamics of an LPS translocon induced by substrate and an antimicrobial peptide. Nature Chemical Biology 17, 187–195 (2020).

64. Deatherage, D. E., & Barrick, J. E. (2014b). Identification of mutations in laboratory-evolved microbes from next-generation sequencing data using breseq. Methods in Molecular Biology, 165–188. 10.1007/978-1-4939-0554-6_12

65. Abueg, L. A., Afgan, E., Allart, O., Awan, A. H., Bacon, W. A., Baker, D., Bassetti, M., Batut, B., Bernt, M., Blankenberg, D., Bombarely, A., Bretaudeau, A., Bromhead, C. J., Burke, M. L., Capon, P. K., Čech, M., Chavero-Díez, M., Chilton, J. M., Collins, T. J., … Zoabi, R. (2024). The Galaxy Platform for Accessible, reproducible, and collaborative data analyses: 2024 update. Nucleic Acids Research, 52(W1). 10.1093/nar/gkae410

66. Pick, K., Stothard, P., & Raivio, T. L. (2024). Complete genome sequence of *Escherichia coli* MP1. Microbiology Resource Announcements, 13(4). 10.1128/mra.01216-23

67. Chen, J., Fruhauf, A., Fan, C., Ponce, J., Ueberheide, B., Bhabha, G., & Ekiert, D. C. (2023). Structure of an endogenous mycobacterial MCE lipid transporter. Nature, 620(7973), 445–452. 10.1038/s41586-023-06366-0

